# Tyramine promotes colon cancer risk and development by inducing DNA damage and inflammation

**DOI:** 10.1101/2023.05.25.542254

**Authors:** Maria Glymenaki, Sophie Curio, Smeeta Shrestha, Mona El-Bahrawy, Yulan Wang, Nigel J. Gooderham, Nadia Guerra, Jia V. Li

**Author notes:** **Corresponding Author**: Jia V. Li, Section of Nutrition, Division of Digestive Diseases, Department of Metabolism, Digestion and Reproduction, Faculty of Medicine, Imperial College London, W12 0NN, London, United Kingdom. The University of Queensland Frazer Institute, The University of Queensland, Woolloongabba, QLD 4102, Australia.

## Abstract

High dietary consumption of processed meat is associated with increased colorectal cancer (CRC) risk, but mechanistic links remain largely unknown. Tyramine is a biogenic amine found in processed food and a gut bacterial product from tyrosine. However, the impact of tyramine on gut health has not been studied. We found that tyramine induced necrosis and promoted cell proliferation and DNA damage in HCT116 cells. Ingestion of tyramine increased colonic tumor size, intestinal cell proliferation and inflammation (e.g., increased mRNA expression of IL-17A and a higher number of Ly6G+ neutrophils) in *Apc^Min/+^* mice. Furthermore, tyramine-treated wild-type mice exhibited visible adenomas and significantly enhanced intestinal tissue DNA damage, together with altered gene pathways involved in epithelial barrier function. In addition, natural killer cell numbers were lower and polymorphonuclear-myeloid derived suppressor cells were higher in tumors from tyramine-treated *Apc^Min/+^* mice, suggesting a suppressive anti-tumor immune response. Thus, tyramine not only increases CRC risk, but also facilitates tumor development. Modulating the levels of tyramine in food and monitoring high-risk individuals could aid in better prognosis and management of CRC.

## Introduction

Colorectal cancer (CRC) is the third most common cancer worldwide and the second cause of cancer-related deaths(Bray *et al*, 2018). Despite a genetic familial origin such as in familial adenomatous polyposis (FAP),(Jasperson *et al*, 2010) environmental factors, such as diet and the gut microbiota, are believed to be important contributors to sporadic CRC. Our previous studies and others reported that high consumption of red or processed meat and fat, and low consumption of fiber could increase CRC risk(Abu-Ghazaleh *et al*, 2021; O’Keefe *et al*, 2015; Song *et al*, 2020). Furthermore, elevated gut bacterial degradation of amino acids and mucins with concomitant depletion of carbohydrate degradation pathways were found in 8 cohorts of patients with CRC(Wirbel *et al*, 2019). Diet is a principal determinant for metabolic function of the gut microbiota; the complex host-diet-gut microbial crosstalk results in a pool of metabolites, which may modulate CRC risk and progression. However, the causal effects of these metabolites on CRC and underlying mechanisms are largely unexplored.

The tyrosine degradation pathway in the gut microbiota was enhanced in patients with CRC(Wirbel *et al*., 2019). A metabolic by-product of dietary tyrosine by bacterial tyrosine decarboxylase is tyramine(Alvarez & Moreno-Arribas, 2014; Bugda Gwilt *et al*, 2019a). Several gut bacteria carry this enzyme, including *Escherichia coli, Enterococcus* and *Klebsiella pneumoniae*(Bugda Gwilt *et al*., 2019a), some of which are abundant in feces and mucosal biopsies from patients with CRC(Dejea *et al*, 2018; Song *et al*., 2020). On the other hand, tyramine, produced by the food-born bacteria, is present in high concentrations in food that are known to increase CRC risk, such as processed meat(Alvarez & Moreno-Arribas, 2014). However, little is known about the function of tyramine apart from its role in increasing blood pressure. A few recent studies showed that tyramine is associated with cytotoxicity and alters the expression profile of genes involved in DNA damage and repair in colonic epithelial cells(Del Rio *et al*, 2018). However, the functional implications of tyramine in the intestinal mucosal health and CRC have not been addressed.

In the current study, we hypothesized that tyramine derived from fermented food and/or gut bacterial metabolism of tyrosine increases CRC risk and promotes CRC development. We tested this hypothesis using both HCT-116 cells, a human colonic epithelial cell line, and *Apc^Min/+^* mice.

## Results

### Tyramine induced cell death and DNA damage

To examine cytotoxic effects of tyramine, HCT116 cells were treated with 0, 0.4, 0.8, 1.2, 1.6, 3.2 or 6.4 mM of tyramine for 24, 48 or 72 hrs. We observed that tyramine was cytotoxic at a concentration of 0.8 mM at 24 hrs, and this cytotoxic effect was persistent up to 72 hrs at the concentration of 1.6 mM or higher (Figure 1A). Cells treated with low concentrations (0.05, 0.15, and 0.2 mM) of tyramine displayed no cytotoxicity (Supplementary Figure 1).

**Figure 1.**
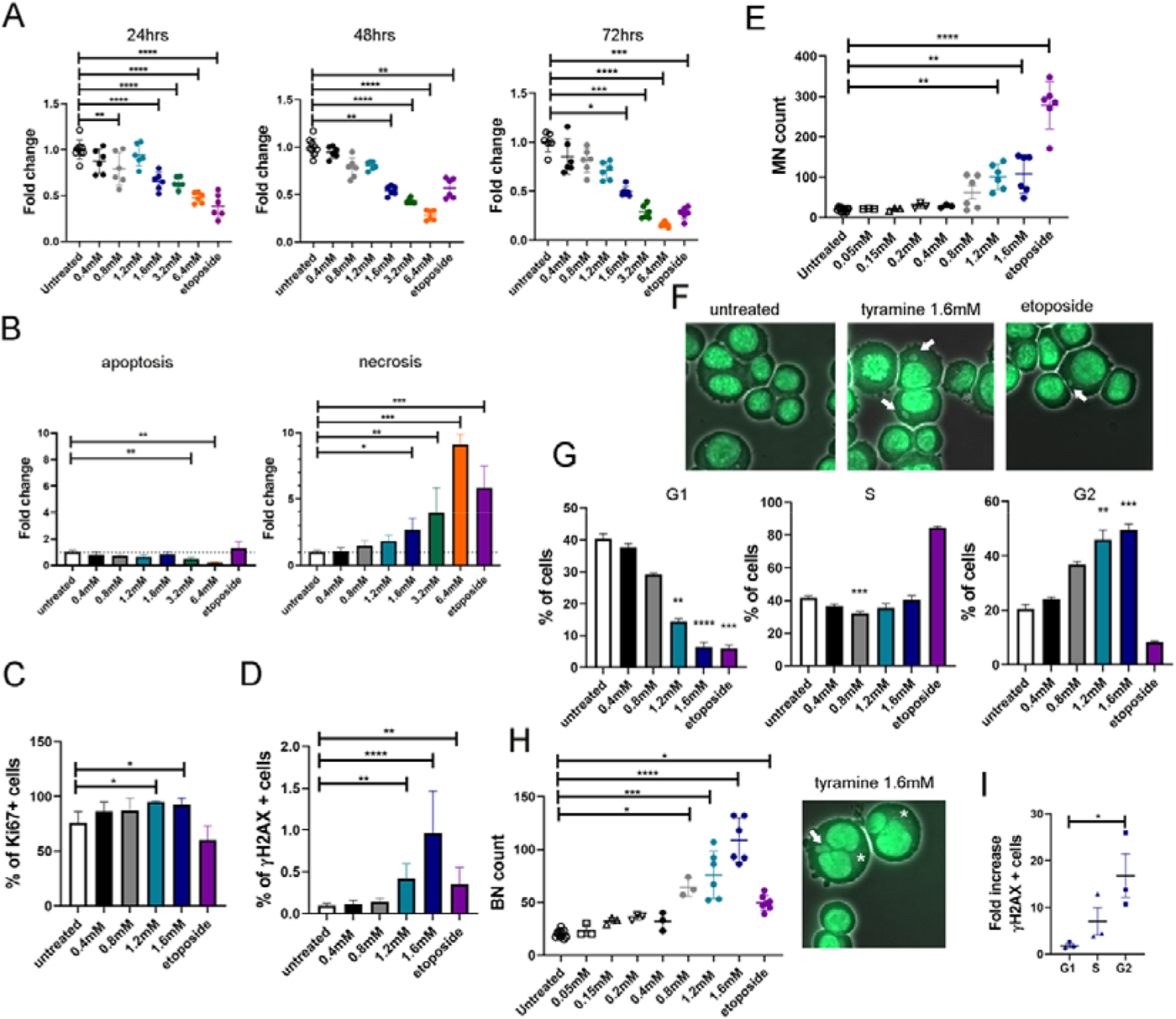
Tyramine reduced cell viability mainly due to necrosis, and increased the number of micronucleated and binucleated cells. A range of concentrations of tyramine was added in human colorectal cancer HCT116 cells over a series of time points. Untreated cells and etoposide served as a negative and positive control respectively for cytotoxicity and genotoxicity. **A)** Fold change of cell viability measured as fluorescence intensity of alamarBlue reagent added to HCT116 cells treated with various concentrations of tyramine for 24, 48 and 72hrs (n=6-9 per group). One-Way ANOVA with Dunnett’s multiple comparisons test was applied for 24hrs. Kruskal-Wallis test with Dunn’s multiple comparisons test for 48 and 72hrs. Results pooled from 3 independent experiments per time point. **B)** Fold change of apoptotic and necrotic cells acquired with flow cytometry using Annexin V and Helix NP Blue DNA dye after 24hrs of treatment with tyramine. Data from two independent experiments were pooled (n=6 per group). Kruskal-Wallis test with Dunn’s multiple comparisons test. **C)** Percentage of Ki67-positive cells after treatment with varying concentrations of tyramine for 24hrs. Data from two independent experiments were combined (n=8 per group). Kruskal-Wallis test with Dunn’s multiple comparisons test. **D)** Percentage of phosphorylated H2AX (γH2AX) positive cells treated with tyramine for 24hrs. Data from two independent experiments were combined (n=8 per group). Kruskal-Wallis test with Dunn’s multiple comparisons test. **E)** Number of micronuclei (MN) per 2000 cells after 48hrs treatment. Slides were scored blindly. Kruskal-Wallis test with Dunn’s multiple comparisons test. **F)** Representative fluorescence microscopy images from MN genotoxicity assay in untreated cells, and cells treated with tyramine 1.6mM and etoposide (40x magnification). Arrowheads show the presence of MN. **G)** Frequency of cells at different stages of cell cycle, namely G1, S and G2/M after 24hr treatment. Kruskal-Wallis test with Dunn’s multiple comparisons test. N=6 per group apart from etoposide group (n=3). **H)** Count of cells with binuclei (BN) per 2000 cells and an image of BN in 1.6mM tyramine on the right. Kruskal-Wallis test with Dunn’s multiple comparisons test. For (**E)** and (**H)** data from three independent experiments were pooled (n=6 for untreated, 1.2mM tyramine, 1.6mM tyramine and etoposide, n=3 for 0.05mM, 0.15mM, 0.2mM, 0.4mM and 0.8mM tyramine). Arrowheads show the presence of MN and asterisks of BN cells. Three slides were scored per treatment. **I)** Fold change of γH2AX positive cells at different stages of cell cycle after 24hr treatment with 1.6mM tyramine over untreated cells. Kruskal-Wallis test with Dunn’s multiple comparisons test. N=3. Data representative from two independent experiments. Data shown as means ± SD for all graphs. *, p<0.05; **, p<0.01, ***, p < 0.001; ****, p < 0.0001.

We further evaluated if tyramine-induced cytotoxicity was contributed by cell apoptosis and/or necrosis by staining live cells with Annexin V and Helix NP Blue dye (Supplementary Figure 2A). The frequency of necrotic cells increased in response to tyramine treatment in a dose-dependent manner (Figure 1B). At high treatment concentrations (3.2 and 6.4 mM), the number of necrotic cells increased with a decrease in apoptotic cell numbers compared to untreated cells (Figure 1B), suggesting a dose-dependent necrosis. In addition, the nuclei without plasma membrane significantly increased with tyramine treatment at a concentration of 1.2 mM or above compared to the untreated control (Supplementary Figure 2B).

Increased cell proliferation and DNA instability are known risk factors for tumorigenesis(Negrini *et al*, 2010). We, therefore, investigated whether tyramine affects cell proliferation and induces DNA damage. The percentage of Ki67 positive cells was significantly higher 24 hrs after addition of tyramine at 1.2 mM and 1.6 mM concentrations compared to the untreated controls (Figure 1C; Supplementary Figure 2C). At these treatment concentrations, a significantly higher frequency of phosphorylated histone H2AX cells, forming γH2AX, was observed in comparison to untreated controls (Figure 1D). This observation indicates a cellular response to the induction of DNA double-strand breaks(Mah *et al*, 2010). Furthermore, we assessed tyramine-induced DNA damage using micronucleus (MN) assay. MN are small extra-nuclear bodies that contain acentric chromosome fragments and/or whole chromosomes. They occur as a consequence of extensive DNA damage, such as double strand breaks and/or deficiencies in chromosome segregation during anaphase(Luzhna *et al*, 2013). Concentrations of tyramine at 1.2 mM and 1.6 mM induced the formation of MN (Figures 1E & 1F), indicating increased DNA damage and chromosomal aberrations. It was not feasible to measure DNA lesions at higher concentrations of tyramine due to high necrosis observed. As we observed that tyramine engendered DNA damage following a 24-hr treatment, we further tested if this genotoxic response is acute by counting γH2AX-positive cells 6 hrs after tyramine treatment at 1.6 mM. Unlike the positive control etoposide that targets directly topoisomerase II generating DNA strand breaks(Baldwin & Osheroff, 2005), tyramine did not induce significant genotoxic effect within 6 hrs of treatment (Supplementary Figure 2D).

Next, we observed that tyramine at 1.2 and 1.6 mM induced cell cycle arrest at the G2/M phase (Figure 1G) and elevated number of binucleated cells (Figure 1H). A significantly higher frequency of γH2AX positive cells following tyramine treatment was found in G2 phase, suggesting the activation of DNA repair mechanisms (Figure 1I). Metabolite profiling of cell media showed separate clustering of tyramine treated and untreated cells suggesting that cell cycle arrest affects cell metabolism and nutrient utilization (Supplementary Figure 3A-C).

It has been shown that pro-inflammatory cytokine, IL-6, regulated cytochrome P450 enzymes (e.g., *CYP1B1*) and enhanced the activation of dietary pro-carcinogens and DNA damage in HCT116 cells(Patel *et al*, 2014). IL-6 also increased invasion and sustained chronic inflammation in co-cultured cancer and immune cells(Patel & Gooderham, 2015). However, our data showed that IL-6 did not exacerbate the cytotoxicity nor genotoxicity induced by tyramine (Supplementary Figure 4A-C; Supplementary Figure 5A-B).

Overall, we showed that tyramine caused cell necrosis, and promoted cell proliferation and DNA damage, which would be expected to compromise genomic stability and increase the risk of colonic tumorigenesis.

### Tyramine affected DNA organization and oxidative respiration pathways

To investigate mechanisms of action of tyramine, we performed RNA-sequencing on untreated control cells and cells treated with 1.2 mM or 1.6 mM of tyramine for 24 hours, which induced significant cytotoxicity and genotoxicity in HCT116 cells. The PCA scores plot of the gene expression profiles displayed a clear separation between control and tyramine-treated cells (Figure 2A). The two tyramine doses induced a similar pattern of cellular responses with a lesser extent from the low dose group. A total of 524 and 241 differentially expressed genes in 1.6 mM and 1.2 mM tyramine-treated cells versus untreated controls were found (Figure 2B and Supplementary Figure 6A). Heatmaps displayed differentially expressed gene sets between tyramine and control groups (Figure 2C and Supplementary Figure 6B). Among the most significantly altered genes were histones (e.g. H1-6, H2AC4, H2BC6, H2BU1, H3C13) which were downregulated, whereas genes involved in oxidation/reduction reactions and oxidative stress response (e.g. AKR1B10, CYP4F3, CYP4F11, OSGIN1) were upregulated in both tyramine-treated groups (Figure 2C). DHRS3 was significantly down-regulated in tyramine-treated groups. Reduced expression of DHRS3 has been previously shown in gastric cancer, and its ectopic expression was found to inhibit cell proliferation and migration(Sumei *et al*, 2021). Expression of genes involved in glucose metabolism such as HKDC1(Zapater *et al*, 2022) and amino acid metabolism such as SLC7A1(You *et al*, 2022), were elevated, suggesting an impact of tyramine on cell energy metabolism.

**Figure 2.**
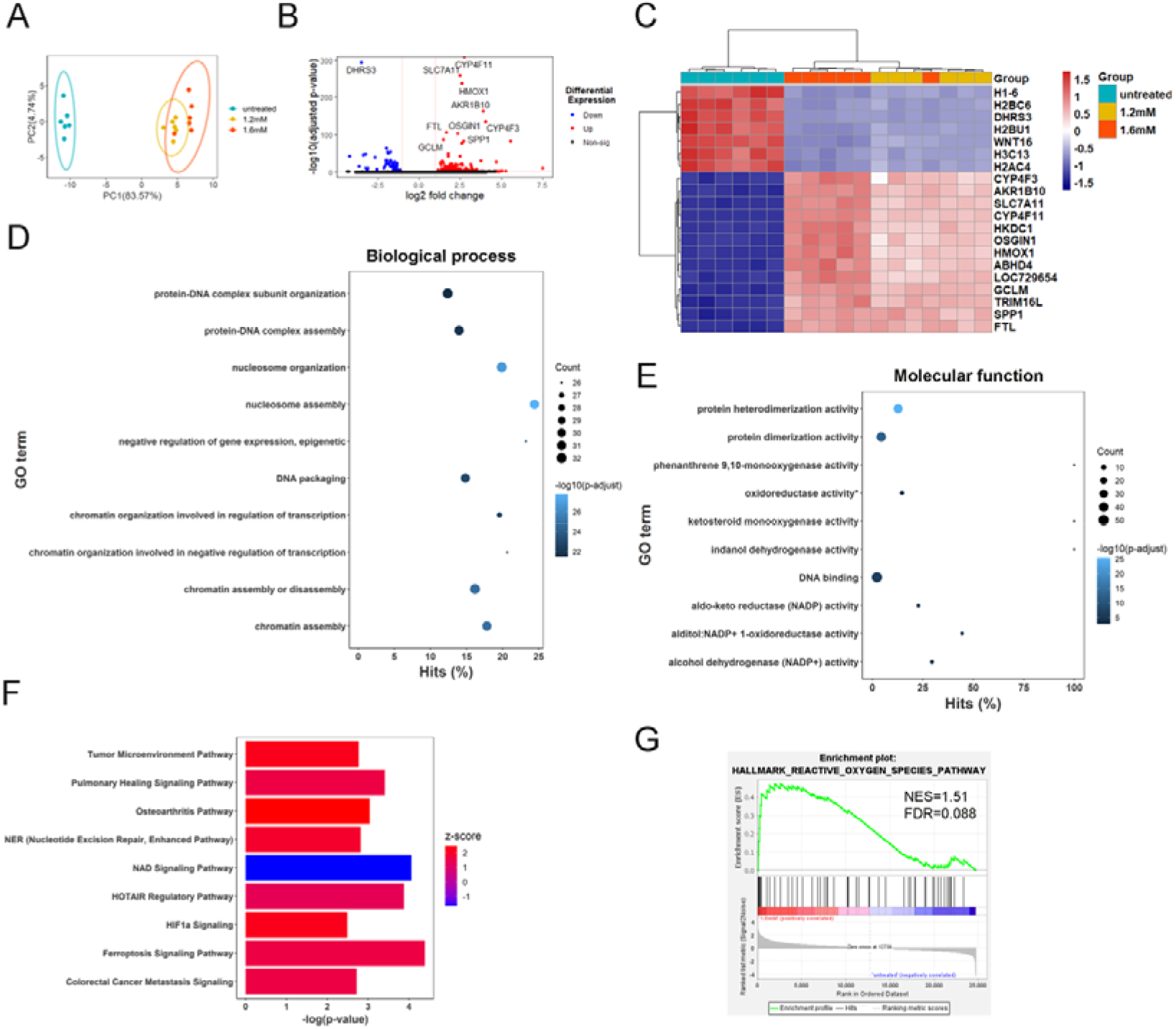
Tyramine alters gene expression profiles and pathways involved in chromatin conformation, oxidative reductive reactions, and colorectal cancer signaling. RNA-seq analysis using DESeq2 on cells treated with 1.2mM or 1.6mM tyramine for 24hrs or left untreated (n=6 per group). **A)** Principal components analysis (PCA) plot showing the clustering of variance stabilizing transformation (VST) RNA-seq data. **B)** Volcano plot of genes enriched or reduced in 1.6mM tyramine treatment versus control. The dotted vertical lines enclose the minimum fold change for the most significant genes. A cut-off of >2 absolute value was used for fold change and <0.05 for FDR-corrected p-value to assign a gene as differentially expressed. The 10 most differentially expressed genes are shown on the plot. **C)** Heatmap of the top 20 genes using z-scores of genes that are differentially expressed between 1.6mM or 1.2mM tyramine treatment group versus control. Each column in the heatmap is an individual sample. **D)** Plot of top 10 over-represented GO terms to visualize the biological processes and **E)** molecular functions altered in 1.6mM tyramine treated cells compared to controls. Hits reflects the proportion of differentially expressed genes in a given pathway. The number of altered genes is indicated by the size of the circle area, and the circle color represents the ranges of the corrected P value. FDR correction < 0.05 has been applied to the depicted GO terms. **F)** IPA pathways that were significantly upregulated (z-score > 0) or downregulated (z-score < 0) in tyramine 1.6mM versus control. Pathways with absolute z-score >1 are depicted. **G)** Gene set enrichment plot demonstrating correlation of gene sets involved in oxygen species pathway with 1.6mM tyramine. Gene permutation was used. FDR correction <0.05 has been applied. NES, normalized enrichment score. Abbreviations: ABHD4, abhydrolase domain containing 4, N-acyl phospholipase B; AKR1B10, aldo-keto reductase family 1 member B10; CYP4F3, cytochrome P450 family 4 subfamily F member 3; CYP4F11, cytochrome P450 family 4 subfamily F member 11; DHRS3, dehydrogenase/reductase 3; FTL, ferritin light chain; GCLM, glutamate-cysteine ligase modifier subunit; HKDC1, hexokinase domain containing 1; HMOX1, heme oxygenase 1; H1-6, H1.6 linker histone, cluster member; H2AC4, H2A clustered histone 4; H2BC6, H2B clustered histone 6; H2BU1, H2B.U histone 1; H3C13, H3 clustered histone 13; LOC729654, inositol polyphosphate multikinase pseudogene; OSGIN1, oxidative stress induced growth inhibitor 1; SLC7A11, solute carrier family 7 member 11; SPP1, secreted phosphoprotein 1; TRIM16L, tripartite motif containing 16 like; WNT16, Wnt family member 16. * denotes oxidoreductase activity, acting on paired donors, with incorporation or reduction of molecular oxygen, NAD(P)H as one donor, and incorporation of one atom of oxygen

Pathway analysis of gene ontology (GO) terms referring to biological process revealed that chromatin organization and assembly networks were significantly reduced in the 1.6 mM tyramine-treated group with an impact on the regulation of transcription and epigenetic control of gene expression (Figure 2D). Consistent with this is the dose-dependent elevated MN frequency (Figure 1E). Chromosomal instability is one of the major routes that leads to CRC and is characterized by structural chromosomal abnormalities, aneuploidy, loss of heterozygosity, and chromosomal rearrangements(Schmitt & Greten, 2021). Therefore, our observation of compromised chromatin assembly and elevated MN frequency strongly indicates signs of chromosomal instability in the presence of tyramine. Furthermore, oxidation and reduction reactions involving NADP/NAD were markedly reduced, suggesting involvement of mitochondria function (Figure 2E and Supplementary Figure 6C). Additionally, IPA analysis consistently showed increased oxidative stress responses and DNA damage repair following tyramine treatment, together with down-regulation of NAD signaling pathway (Figure 2F). Moreover, processes related to colorectal cancer metastasis signaling and tumor microenvironment were enhanced. Ferroptotic cell death, which is iron-dependent and results from the accumulation of lipid reactive oxygen species (ROS)(Li *et al*, 2020a), was higher in the tyramine group. In further support of the role of tyramine in oxidative damage, Gene Set Enrichment Analysis (GSEA) showed overrepresentation of gene sets in reactive oxygen species (Figure 2G and Supplementary Figure 6D). Analogous findings of gene pathway analysis were also found in the 1.2 mM tyramine group (Supplementary Figure 7A-F). The RNA sequencing data analyses collectively indicate that tyramine impacted DNA structure and resulted in oxidative stress, which may be responsible for its genotoxic and cytotoxic effects.

### Tyramine facilitated tumorigenesis in *Apc^Min/+^* mice and increased CRC risk in WT mice

Following up on the observation of tyramine-induced cyto- and geno-toxicity *in vitro*, we used *Apc^Min/+^* mice to examine the impact of tyramine in CRC development *in vivo* (Figure 3A). No significant differences in body weight, food consumption and water intake were observed between any of the experimental groups throughout the study (Figures 3B-D). Spleen weight was significantly higher, and hematocrit was significantly lower in *Apc^Min/+^* mice compared to wild-type controls irrespective of the treatment regime (Figure 3E and F). These findings mirror previous results(Chae & Bothwell, 2011; Myzak *et al*, 2006) reported in *Apc^Min/+^* mice that develop anemia concomitant with tumor growth(Li *et al*, 2012). In contrast, the liver weight was comparable among the groups (Figure 3G). While the total gut length was comparable between tyramine-treated and untreated groups, a significantly shorter gut length was noted in tyramine-treated *Apc^Min/+^* mice compared to tyramine-treated WT mice (Figure 3H), which suggests a crosstalk between tyramine and genetic strains of mice resulting in this phenotypical change in the gut length. Colon length was similar among the groups (Supplementary Figure 8A). Histological analysis indicated a similar grade of dysplasia in the CRC-prone mice independent of treatment, and tyramine administration did not result in histological changes (Figure 3I).

**Figure 3.**
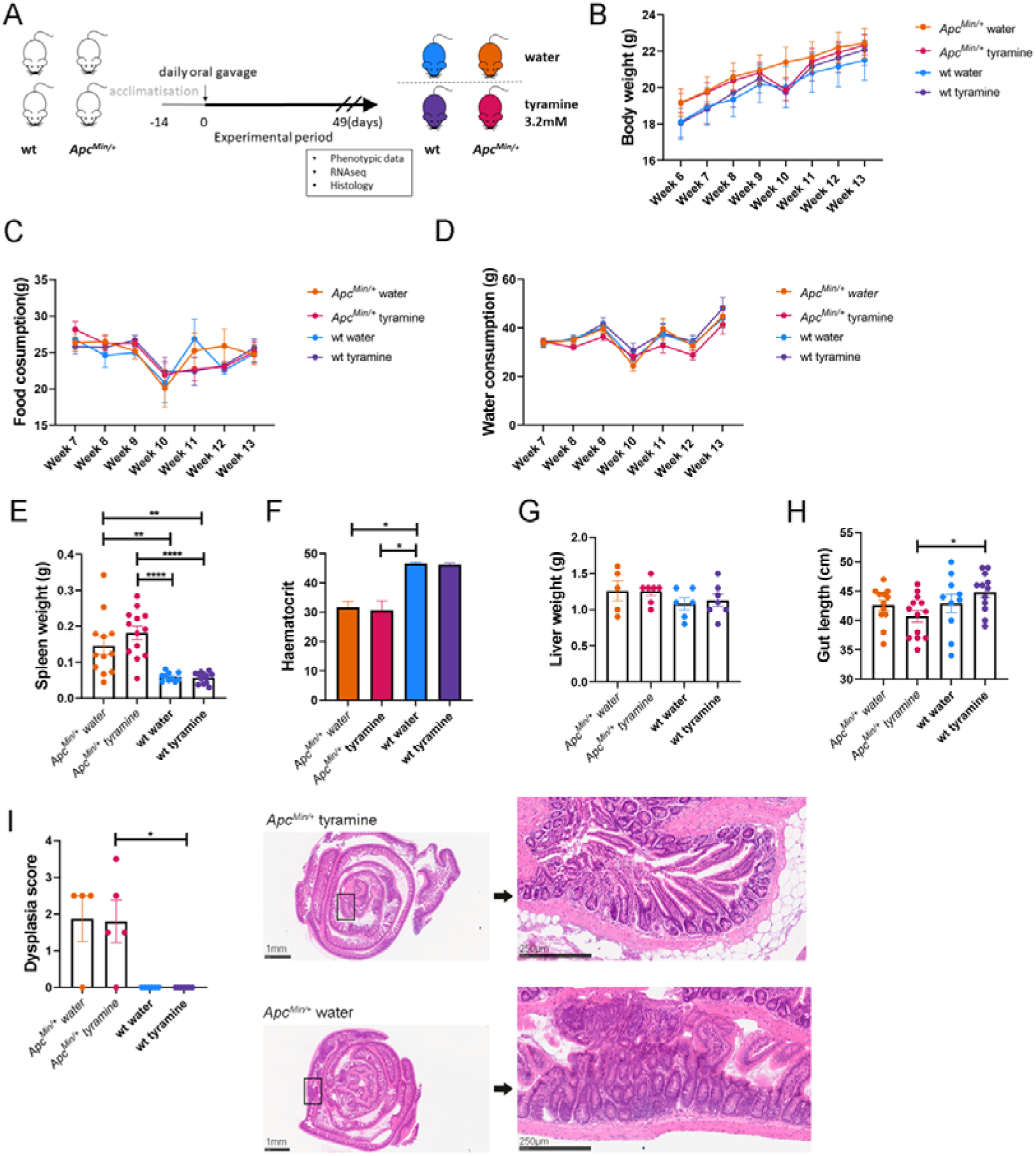
Phenotypic characterization of tyramine-treated *Apc^Min/+^* and control WT mice. *Apc^Min/+^* mice and respective WT littermate controls received daily doses of 3.2mM tyramine (n=13 for *Apc^Min/+^* mice and n=12 for WT mice) or water (n=12 for *Apc^Min/+^* mice and n=10 for WT mice) via oral gavage for a period of 49 days. Treatment started at 6 weeks of age and concluded at 13 weeks of age. **A)** Schematic of the study design. **B)** Body weight, **C)** Food consumption and **D)** Water intake across the study among tyramine-treated *Apc^Min/+^* mice, untreated *Apc^Min/+^* mice, tyramine-treated WT and untreated WT mice. **E)** Spleen weight, **F)** Hematocrit, **G)** Liver weight and **H)** Gut length at 13 weeks of age. **I)** Dysplasia scoring based on ileum gut rolls taken at study endpoint and representative H&E sections from tyramine-treated and untreated *Apc^Min/+^* mice. Scale bar 1mm for whole gut roll image. Square indicates the area of magnified image. Scale bar 250µm for magnified image. Data shown as means ± SEM. Two-way ANOVA with Tukey’s multiple comparisons test in B), mixed effects model in C) and D), ordinary one-way ANOVA with Tukey’s multiple comparisons test in E) and H), Kruskal-Wallis test with Dunn’s multiple comparisons test in F), G) and I). *, p<0.05; **, p<0.01, ****, p < 0.0001.

Macroscopic evaluation showed that *Apc^Min/+^* mice receiving tyramine tended to develop a higher number of tumors, especially in the colon (Figure 4A). The number of mice and the percentages of mice with tumors were consistently higher in the tyramine-treated *Apc^Min/+^* group compared with the water-treated *Apc^Min/+^* group across the GI tract (Figure 4B and Supplementary Figure 8B). In WT mice where tumors were not expected, tyramine treatment resulted in the development of visible adenomas in 5 out of 12 mice in contrast to none in water-treated WT mice. The number of tumors displayed a positive correlation with spleen weight (r = 0.79, p= 2.7e-11; Supplementary Figure 8C). Appearance scoring demonstrated that tyramine administration tended to exacerbate the pathology in *Apc^Min/+^* mice (Supplementary Figure 8D). We found that the total number of tumors was higher but not statistically significant in tyramine-treated *Apc^Min/+^* mice compared to untreated *Apc^Min/+^*mice (Figure 4C-E, Supplementary Figure 8E). However, tyramine had a significant effect on the size of the tumors in the colon (Figure 4F). The diameters of intestinal adenomas in mice were measured and allocated to either one of three categories: diameter <1 mm, 1–3 mm, and >3 mm. Tumors with size 1-3 mm were in greater abundance in the colon of tyramine-treated *Apc^Min/+^* mice compared with untreated *Apc^Min/+^* controls (Figure 4F).

**Figure 4.**
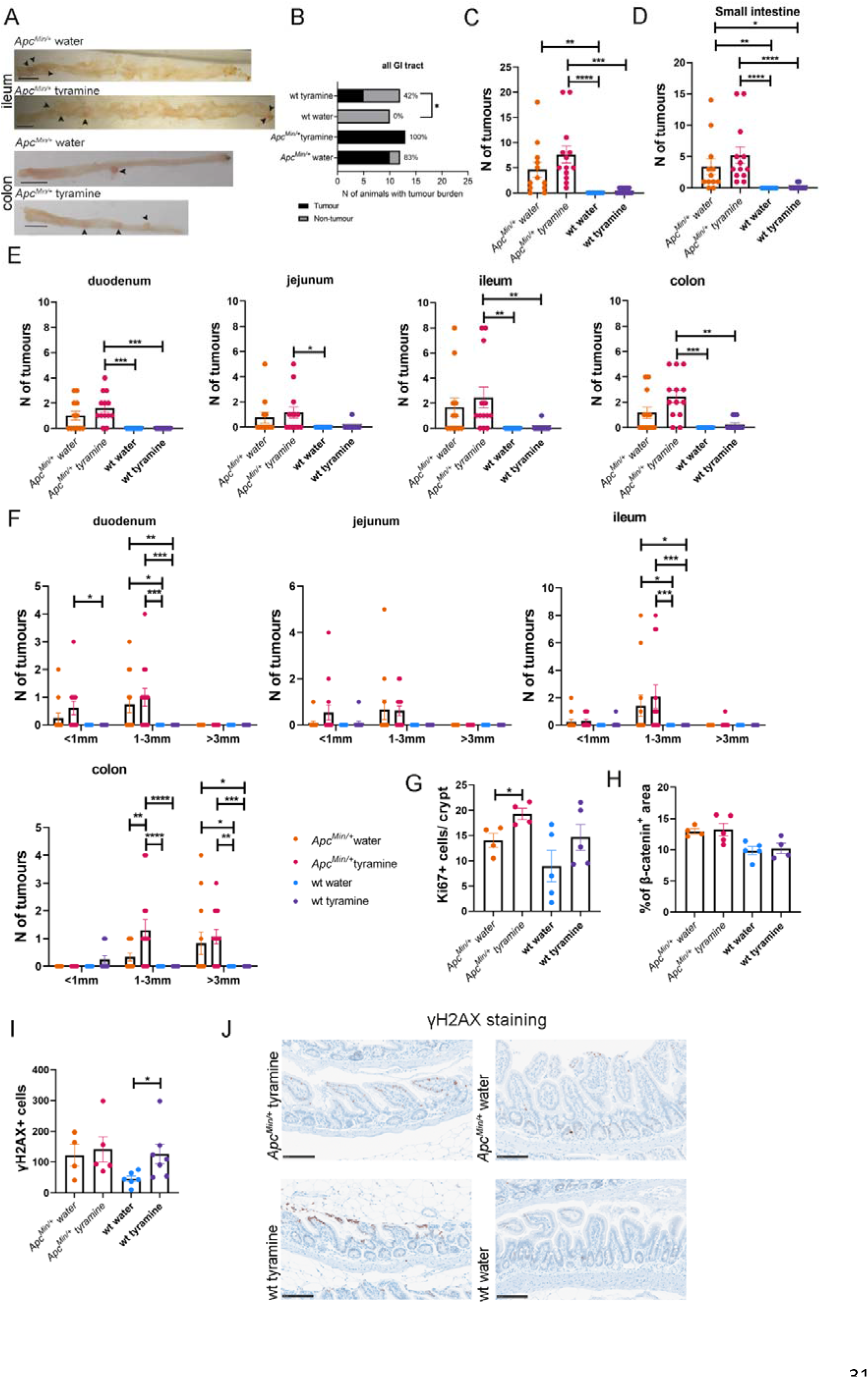
Tyramine promotes CRC development in *Apc^Min/+^* mice and increases CRC risk in WT controls. **A)** Representative macroscopic images of ileum and colon in *Apc^Min/+^* mice administered water or tyramine. Scale bars, 1 cm. **B)** Contingency table showing the number of animals with tumors. The percentages of animals having tumors per group is also depicted. **C)** Total tumor numbers of the small and large intestine were counted in 13-week old tyramine-treated *Apc^Min/+^* mice (n=13), untreated *Apc^Min/+^* mice (n=12), tyramine-treated WT mice (n=12) and untreated WT mice (n=10). **D)** Small intestine tumors, and **E)** tumors numbers from each part of the small intestine (duodenum, jejunum, ileum) and colon. **F)** Tumor numbers based on tumor size. Tumors smaller than 1 mm, between 1–3 mm, and greater than 3 mm were counted separately. **G)** Number of Ki67+ cells per crypt in ileum. A total of 60 crypts per slide was counted. 3 slides were acquired for each tissue sample. **H)** Quantification of β-catenin positive area. **I)** Number of γH2AX positive cells per field of view. 3 separate fields of view were taken for each tissue sample. **J)** Representative γH2AX immunohistochemistry images in ileum. Scale bar 100 µm. Data shown as means ± SEM. Fisher’s exact test in B) and Kruskal-Wallis test with Dunn’s multiple comparisons test in C) – E). Two-way ANOVA with Tukey’s multiple comparisons test in F). Mann-Whitney test for pairwise comparisons between tyramine-treated mice and water-treated *Apc^Min/+^*mice, and tyramine-treated and water-treated WT mice in G), H) and I). *, p<0.05; **, p<0.01, ***, p < 0.001; ****, p < 0.0001.

Tyramine treatment resulted in a higher number of Ki67+ cells when comparing within the same genotype, with a statistically significant change in *Apc^Min/+^* mice (Figure 4G, Supplementary Figure 9A). In contrast, β-catenin levels were similar among the groups (Figure 4H and Supplementary Figure 9B). These data indicate that tyramine treatment increases cell proliferation without affecting the Wnt/ β-catenin signaling pathway. Furthermore, while no difference in the number of γH2AX+ cells was noted between tyramine-treated and untreated *Apc^Min/+^* mice, a significantly enhanced number of γH2AX+ cells was observed in tyramine-treated WT mice compared to the control, indicating that tyramine induced DNA damage (Figure 4I-J). These data collectively showed that tyramine was tumor-promoting in the APC-model of intestinal tumorigenesis and promoted DNA-damage in WT mice.

### Tyramine impacted gene pathways involved in inflammation and extracellular matrix in the ileum and colon of *Apc^Min/+^* mice

Tyramine binds to a family of G-protein-coupled receptors, termed trace amine-associated receptors (TAARs), which are expressed in the intestine(Bugda Gwilt *et al*., 2019a). There is currently no information on TAAR1-mediated uptake of tyramine in gut epithelial cells, but it has been shown that tyramine can be absorbed through diffusion(Tchercansky *et al*, 1994). We did not detect tyramine in urine or feces when administered orally (Supplementary Figure 10), suggesting that it is absorbed. To gain a deeper understanding into its function in tumorigenesis *in vivo*, we performed transcriptomic analysis of tumor-free colon and ileum tissues of *Apc^Min/+^* and WT mice. We compared transcriptional profiles between tyramine- and water-treated mice in each genotype to focus on the effect of treatment.

Transcriptomic analyses of both ileum and colon tissue from *Apc^Min/+^*mice showed upregulated expression levels of genes involved in inflammatory responses to tyramine. Colonic tissue expression of *IL-17A* gene was found to be up-regulated in the tyramine-treated compared to untreated *Apc^Min/+^*mice (Figure 5A). IL-17A is a pro-inflammatory cytokine that drives intestinal tumorigenesis(Hurtado *et al*, 2018) and it is required for tumor formation in *Apc^Min/+^* mice(Chae *et al*, 2010). In addition, we observed higher expression levels of neutrophilic granule protein (*Ngp*) gene in the ileum tissue of tyramine-treated *Apc^Min/+^* mice in contrast to untreated *Apc^Min/+^* mice (Figure 5B). This observation is consistent with the immunohistochemistry analysis of the ileum tissue, which showed a higher number of Ly6G+ cells (neutrophils) in *Apc^Min/+^* mice following tyramine treatment compared to untreated controls (Supplementary Figure 11A-B).

**Figure 5.**
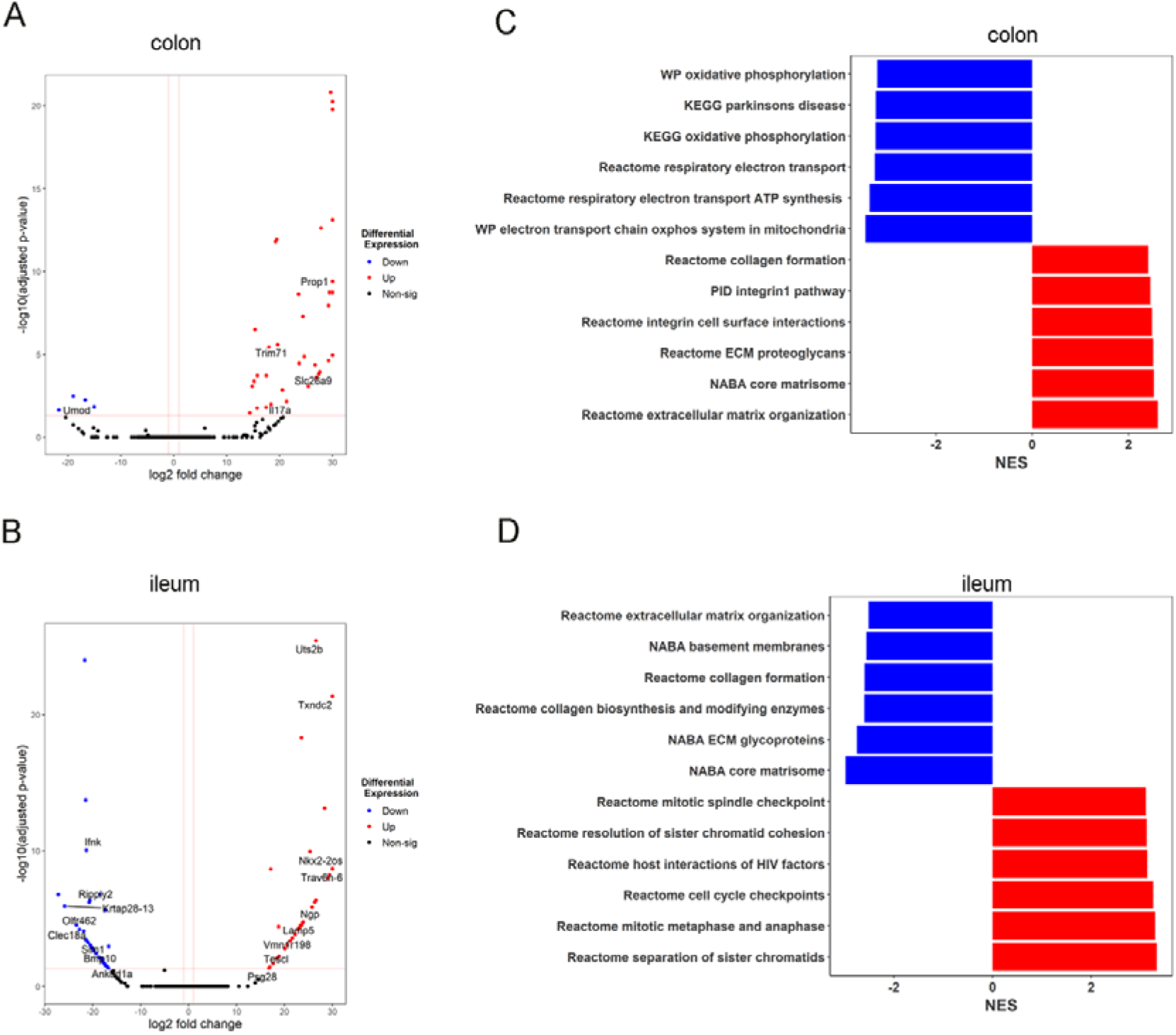
Tyramine promotes an inflammatory environment in surrounding healthy gut tissue instigating tumor initiation in *Apc^Min/+^* mice. RNA-seq analysis using DESeq2 on colon and ileum tissue of *Apc^Min/+^* mice treated with tyramine 3.2mM or water (n=6 for colon tyramine-treated *Apc^Min/+^* mice, n=5 for colon water-treated *Apc^Min/+^* mice, and n=10 per each group for ileum). Volcano plot of genes enriched or reduced in tyramine treatment versus control in **A)** colon and **B)** ileum. The dotted vertical lines enclose the minimum fold change for the most significant genes. A cut-off of >2 absolute value was used for fold change and <0.05 for FDR-corrected p-value to assign a gene as differentially expressed. Differentially expressed genes are shown on the plot. Abbreviations of some interesting genes: *IL17a*, interleukin -17A; *Ngp*, neutrophilic granule protein. **C)** GSEA of the top-6 most significant downregulated and upregulated pathways in colon and **D)** ileum. FDR correction <0.05 has been applied. NES, normalized enrichment score.

GSEA pathway analysis showed enrichment of genes involved in extracellular matrix organization and decreases in mitochondrial activities in the colon tissue following tyramine treatment (Figure 5C). In contrast, extracellular matrix organization pathway was downregulated in the ileum, together with collagen biosynthesis pathways in the tyramine-treated *Apc^Min/+^* mice compared to untreated ones, which may indicate changes in gut epithelial barrier (Figure 5D). The different observations in the colon and ileum could be attributed to their physiological differences and possibly differential binding of tyramine to its relevant receptors across the intestine(Bugda Gwilt *et al*., 2019a). In the tumors, pathways including extracellular matrix glycoproteins and basement membranes (BM) were down-regulated by tyramine treatment, which may indicate an epithelial-mesenchymal transition (EMT) in the tumors and invasion to the surrounding stroma through BM(Elgundi *et al*, 2019) (Supplementary Figure 11C).

### Tyramine impacted the cell cycle and epithelial barrier function pathways in the ileum and colon of WT mice

We found a significant upregulation of the cell division cycle gene (*Cdc34b*) expression in the colon of tyramine-treated WT mice compared to water-treated mice (Figure 6A). Moreover, the expression levels of genes related to epithelial barrier function, including defensins and C-type lectins (*Defb14* and *Clec2f*) were elevated in the colon and ileum, respectively, in the tyramine-treated WT mice compared to water-treated WT mice (Figure 6B). Similar to tyramine-treated *Apc^Min/+^* mice, pathways related to extracellular matrix components were overrepresented in the colon of the tyramine-treated WT mice, whereas cell cycle control from G1 to S phase was diminished, suggesting reduced control in cell proliferation (Figure 6C). Protein translation and synthesis pathways (i.e. Reactome eukaryotic translation elongation, Reactome SRP depended cotranslational protein targeting to membrane, KEGG ribosome, Reactome regulation of expression of slits and robos, Reactome rRNA processing, Reactome translation) were upregulated in the tyramine-administered group in the ileum, suggesting increased cell division (Figure 6D). These observations are consistent with higher levels of Ki67+ cells in tyramine-treated WT mice and tyramine-treated HCT116 cells in contrast to their untreated controls.

**Figure 6.**
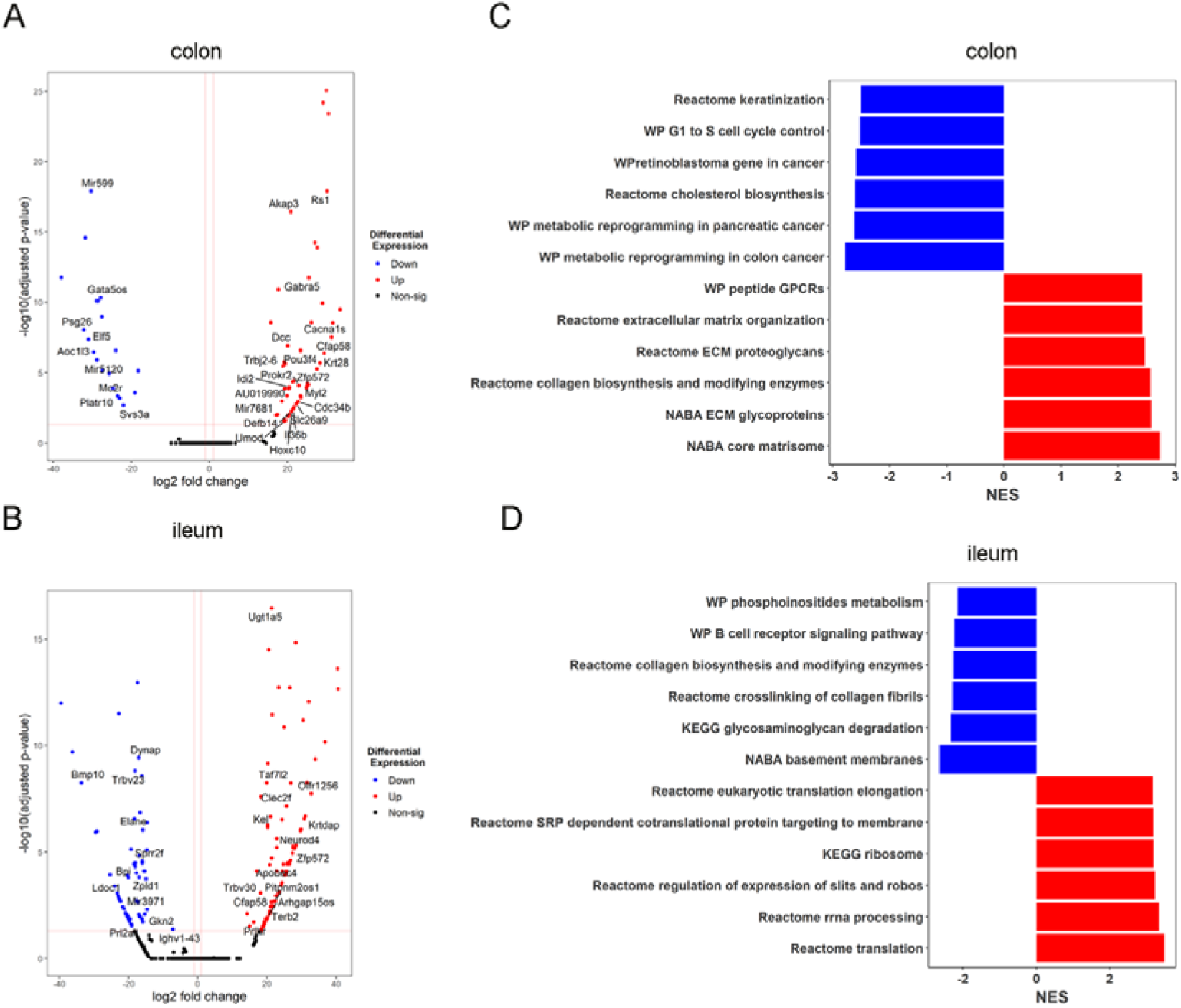
Tyramine alters the expression profiles of genes involved in epithelial barrier function in wild-type mice. RNA-seq analysis using DESeq2 on colon and ileum tissue of WT mice treated with tyramine 3.2mM or water for 7 weeks (n=5 for colon tyramine-treated WT mice, n=4 for colon water-treated WT mice, and n=9 for ileum tyramine-treated WT mice, n=8 for ileum water-treated WT mice). Volcano plot of genes enriched or reduced in tyramine treatment versus control in **A)** colon and **B)** ileum. The dotted vertical lines enclose the minimum fold change for the most significant genes. A cut-off of >2 absolute value was used for fold change and <0.05 for FDR-corrected p-value to assign a gene as differentially expressed. Differentially expressed genes are shown on the plots. Abbreviations of some interesting genes: *Cdc34b*, cell division cycle 34B; *Defb14*, defensin beta 14; *Clec2f*, C-type lectin domain family 2, member f. **C)** GSEA of the top-6 most significant downregulated and upregulated pathways in colon and **D)** ileum. FDR correction <0.05 has been applied. NES, normalized enrichment score.

Collectively, our data suggest that tyramine drives an inflammatory environment in the healthy gut tissue as evidenced by the upregulation of *IL-17A* and *Ngp* genes in *Apc^Min/+^* mice and is associated with increased cell proliferation and impaired epithelial barrier in the *Apc-* deficient and wild-type settings.

### Tyramine promotes a tumorigenic immune response in tumor microenvironment

Tyramine has been shown to increase inflammatory gene expression in macrophages(Bugda Gwilt *et al*, 2019b). Therefore, we further investigated functions of the immune cell subsets within the ileal and colonic tumors using flow cytometry (Supplementary Figure 12A-B). Due to large variation in tumor numbers and size from the *Apc^Min/+^* mice, we therefore collected tumor samples from *Apc^Min/+^* mice (3 control and 4 tyramine-treated mice) for flow cytometry analysis. We observed similar frequencies on lymphoid, innate lymphoid and myeloid cells with the exception of NK cells and polymorphonuclear-myeloid derived suppressor cells (PMN-MDSCs) in colonic tumor infiltrating lymphocytes (TIL) (Figure 7A; Supplementary Figure 13 and Figure 14A). The frequency of NK cells was lower in tyramine-treated compared to water-treated *Apc^Min/+^* mice. This was accompanied by a reduction in the mean fluorescence intensity (MFI) of NKG2D in NK cells in the tumor microenvironment (Figure 7B). NK cells, in particular NKG2D-expressing NK cells, are important players in the anti-tumor response due to their ability to recognize and eliminate tumor cells without the need for antigen-specific receptors(Chiossone *et al*, 2018; Dhar & Wu, 2018). Our observation of a reduction in NKG2D expressing NK cells suggested that tyramine may compromise an effective anti-tumor response at the early stage of tumorigenesis. On the other hand, PMN-MDSCs displayed an increased trend in their proportion in the tyramine-treated group (Figure 7A). PMN-MDSCs have several functions but are generally thought to suppress the anti-tumor response, thereby promoting tumor growth(Gabrilovich *et al*, 2012). Furthermore, MFI of the inhibitory marker PD-1 showed a higher frequency in CD8+ T cells but not reaching statistical significance (Figure 7C), suggesting previous activation and potential exhaustion of cytotoxic CD8+ T cells. Tumor-specific CD8+ T cells become exhausted due to persistent stimulation impairing their effector function against tumor progression(Jiang *et al*, 2020). We did not observe any statistically significant differences in the functional immune markers that we measured in CD4+ T cells, γδT cell, innate lymphoid cell 1 (ILC1) and ILC3 (Supplementary Figure 14B). However, it is worth noting that we observed clear lower percentages of IL-17 expressing CD4+ T cells, and TNF-α expressing CD4+ and CD8+ T cells and NK cells in colonic tumors from tyramine-treated mice compared to water-treated mice. In contrast, GzB expressing γδT cells were increased following tyramine treatment. Our findings suggest that tyramine suppresses an anti-tumor immune response to favor tumor progression.

**Figure 7.**
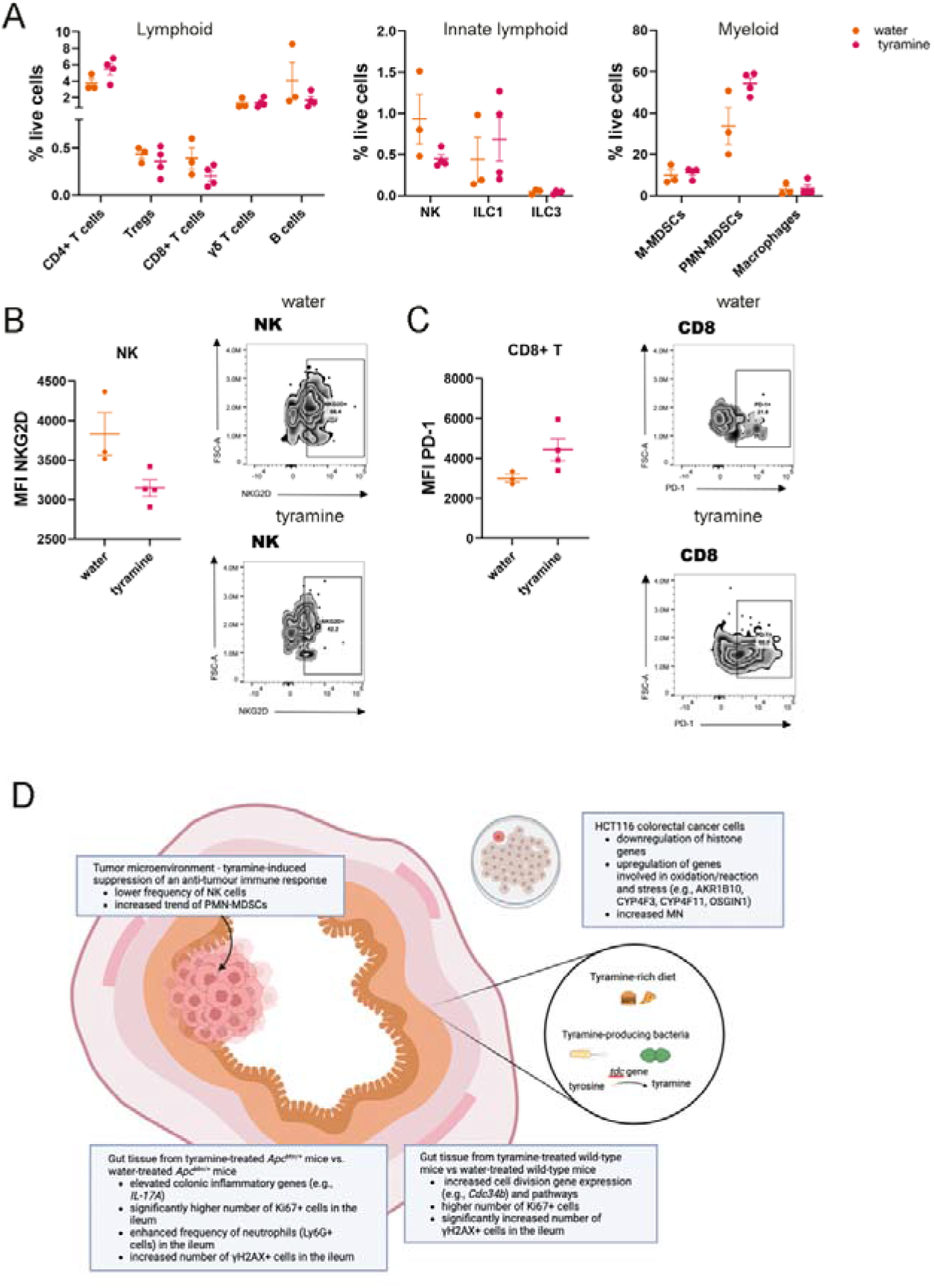
Tyramine dampens an anti-tumor immune response in colonic tumors. **A)** Lymphoid, innate lymphoid and myeloid tumor immune cell infiltrates plotted as percentage of live non-autofluorescent cells in tyramine- and water-treated *Apc^Min/+^*mice at 13 weeks of age. **B)** Mean Fluorescence Intensity (MFI) of NKG2D in NK cells. Representative flow cytometry zebra plots for tyramine- and water-treated mice are also shown. **C)** MFI of PD-1 in CD8+ T cells and representative zebra plots. Data represent mean ± SEM of two independent experiments (N=3-4 per group) in **A-C**. Wilcoxon Rank Sum test was applied for between group comparisons followed by fdr method for adjusting p-values for multiple test correction **(A-C)**. **D)** Schematic on the effect of tyramine in CRC development.

## Discussion

We reported that tyramine was cytotoxic and genotoxic to HCT116 cells, and induced cell cycle arrest and necrosis. Expression levels of genes involved in chromosome organization and oxidative stress were upregulated in tyramine-treated HCT116 cells. To the best of our knowledge, the study herein is the first one addressing the role of tyramine in CRC development *in vivo*. The dose of tyramine we used here is equivalent to 220 mg intake for a person with a body weight of 50 kg, which mirrors the human consumption of foodstuff high in tyramine(Andersen *et al*, 2019; Linseisen *et al*, 2006; Paulsen *et al*, 2012). Our experiment using *Apc^Min/+^* mice showed that tyramine significantly exacerbated tumorigenesis, induced pro-inflammatory responses and increased cell proliferation (Figure 7D). Most importantly, tyramine induced a significantly higher level of DNA damage in the colon of the WT mice, accompanied by a trend for increased cell proliferation and tumorigenesis was observed in a small fraction of WT mice treated with tyramine.

Our observation of tyramine-induced cytotoxicity is consistent with previous studies, where tyramine was tested at higher concentrations (e.g., 14.58 mM) using HT29 human colonic cells(Del Rio *et al*, 2017; Linares *et al*, 2016). In our study, tyramine started to induce significant cytotoxic effects at 0.8 mM. These differences in cytotoxic doses could be related to different cell lines used in these experiments. Moreover, we identified necrosis as the main form of cell death 24-hr post tyramine treatment, consistent with previous findings in HT29 cells(Linares *et al*., 2016). In contrast, apoptosis-involved genes, such as Caspase 7 (CASP7) and BCL2 Binding Component 3 (BBC3) were reported to upregulate in HT29 cells following a 6-hr tyramine treatment at 14.58 mM(Del Rio *et al*., 2018). These data may suggest apoptosis is an acute cell response to tyramine and it shifts to necrosis as time and concentrations increase, as supported by our data in which apoptosis alone was significantly reduced at higher treatment concentrations.

Genomic instability is a risk factor for cancer(Negrini *et al*., 2010). Our data showed that tyramine increased the frequency of γH2AX+ cells and promoted the frequency of MN formation in tyramine-treated HCT116 cells. Furthermore, the number of γH2AX+ cells was significantly higher in the colon of tyramine-treated WT mice compared to water-treated WT mice. These observations indicate that tyramine can induce DNA damage, suggesting its role in increasing CRC risk. In agreement with our findings, altered expression levels of genes involved in DNA damage and repair pathways such as ERCC3, GADD45G, POLB, and PPP1R15A were reported in HT29 cells treated with tyramine(Del Rio *et al*., 2018). Furthermore, we found that tyramine downregulated the expression of histones, whereas genes involved in oxidation/reduction reactions and oxidative stress response (e.g., AKR1B10, CYP4F3, CYP4F11, OSGIN1) were upregulated. Pathways involved in chromatin organization and assembly were reduced, suggesting genomic instability in response to tyramine treatment. In addition, we observed that tyramine induced cell cycle arrest at G2, when DNA damage occurred. It is likely that cells undergo an arrest of the cell cycle allowing for DNA repair to occur(Scully *et al*, 2019). We also found an increase in the percentage of Ki67 positive cells and impaired mitochondria-related functions after tyramine treatment *in vitro* and *in vivo*. It has been reported that biological pathways associated with DNA damage repair and proliferation were enhanced at early stages of CRC, whereas processes related to mitochondrial function were reduced (bioRxiv 2021.09.30.462502). With the additional observation of the development of adenoma formation in WT mice receiving tyramine, these data collectively prompt us to conclude that tyramine increases CRC risk by increasing DNA damage and cell proliferation.

Another observation was that tyramine increased tumor size in the colon and exacerbated the appearance of *Apc^Min/+^*mice compared to water-treated mice, suggesting that it facilitates tumor development. A previous study on the metabolic modeling of the gut microbiome identified tyramine as a significant metabolite in CRC microbiome(Salahshouri *et al*, 2021). These evidence support the hypothesis that tyramine derived from the diet and/or microbial metabolism facilitates CRC development, particularly under circumstances where there is a genetic predisposition as is the case with a mutated APC gene. It is interesting to speculate whether a similar situation exists with other CRC genetic predispositions such as mutant RAS or p53. Furthermore, RNA-seq analysis revealed that tyramine impacted cell division and disrupted the extracellular matrix organization in *Apc^Min/+^* and wild-type mice. Extracellular matrix composition and structure are deregulated in CRC to promote tumor growth(Li *et al*, 2020b). Changes in extracellular matrix promoted the proliferation of cancer cells(Li *et al*., 2020b). Enrichment in extracellular matrix organization pathways has also been previously reported in a colitis-associated model of CRC indicating their importance for tumor initiation(Hammad *et al*, 2021).

Another profound finding was that tyramine increased the levels of *IL-17A* gene expression in the colon of *Apc^Min/+^* mice compared to the water-treated mice. Furthermore, *Ngp* gene expression was higher in the ileum of tyramine-treated *Apc^Min/+^* mice. The Ngp protein belongs to the cystatin family with a critical role in the regulation of inflammatory response(Liu *et al*, 2020). Ly6G+ cells identified as neutrophils were in greater abundance in the ileum of *Apc^Min/+^* mice treated with tyramine compared to water. These data suggest that tyramine promotes an inflammatory environment in the gut mucosa. Innate immune pathways driving inflammation are increased during CRC onset in the epithelium (bioRxiv 2021.09.30.462502). Loss of *Apc* induces disruption of epithelial barrier activating tumor-associated myeloid cells by components of the gut microbiota, which regulate IL-17 production(Grivennikov *et al*, 2012). Our data suggest that tyramine contributes to epithelial barrier impairment and drives an inflammatory response. In addition, NK cells were lower in tumors in tyramine-treated *Apc^Min/+^* mice compared to water-treated *Apc^Min/+^* mice, while PMN-MDSCs displayed an increasing trend. These observations suggest that tyramine has a potential to suppress an effective anti-tumor immune response, favoring tumor progression. The reduction in the proportion of NK cells in tumors of tyramine-treated *Apc^Min/+^* mice may result in reduced lysis of tumor cells, hence increasing tumor size. These immune responses are consistent with a previously reported transcriptomic analysis of CRC patients, showing that anti-tumor immune responses including activation of NK cells and leukocyte mediated cytotoxicity were downregulated in the initial stages of normal to adenoma transition(Hong *et al*, 2021). Previous studies have shown that NK cells were rarely detected in colonic tumors from *Apc^Min/+^* mice(Li *et al*., 2012), whereas PMN-MDSCs accumulation inhibited anti-tumor T-cell responses(Chun *et al*, 2015). These data collectively indicate that tyramine could contribute to tumor development through an induction of pro-inflammatory response and suppression of anti-tumor immune responses.

In summary, our data highlight the function of tyramine in increasing CRC risk and promoting tumorigenesis *in vivo*. These findings provide insight into nutritional prevention to reduce risk of CRC. This study demonstrates an example that exposure to certain dietary and microbial metabolites could affect tumor progression and modulating these metabolite levels could improve CRC prognosis.

## Materials and Methods

### Cell culture

The HCT116 cell line, a human colorectal adenocarcinoma adherent cell line, was purchased from the American Type Culture Collection (ATCC, RRID: CVCL_0291, LGC Prochem). Cells were routinely cultured in RPMI 1640 medium (Gibco) supplemented with 10% Fetal Bovine Serum (FBS), 100 units/mL penicillin, 100 μg/mL streptomycin, and 2 mmol/L L-glutamine (Gibco). The cells were passaged every 2-3 days and passages 3-7 were used in the current study.

### Cell treatment

Cells were maintained in culture medium supplemented with 5% dextran-coated charcoal-stripped FBS for 72 hrs to remove hormones and growth factors before any treatment. The cells were seeded and treated with various concentrations of tyramine dissolved in water (0.2 mM to 6.4 mM, pH=7.01) for 24, 48 or 72 hrs. Specifically, 20,000 cells per well were used for cell viability, 40,000 for genotoxicity and 100,000 cells for flow cytometry assays. To assess the impact of inflammatory environment on tyramine treatment, cells were treated with interleukin-6 (IL-6, Sigma) for 24, 48 and 72 hrs at a range of 0-1000 pg/ml as previously reported(Patel & Gooderham, 2015). Micronucleus assay and cell staining for apoptosis, cell cycle, proliferation and DNA damage analysis using flow cytometry are described in Supplementary Information.

### Animal experiment design

Mouse model and breeding are described in the Supplementary Information. *Apc^Min/+^* and wild-type (WT) mice were kept under specific pathogen-free conditions at the Central Biomedical Services facility at Imperial College London in a 12-hour light/dark circle. The health status of mice was routinely monitored throughout the study. The experiment was performed in accordance with the Animals (Scientific Procedures) Act 1986 under the Project License (9718F9C8) issued by the UK Home Office.

*Apc^Min/+^* and WT mice at 6 weeks of age were oral gavaged daily with 200 µL of sterile-filtered tyramine solution at 3.2 mM and pH=7.01 (Sigma-Aldrich) or water for 7 weeks. The dosage was determined based on dietary intake of foods that contain high amounts of tyramine in humans(Andersen *et al*., 2019; Linseisen *et al*., 2006; Paulsen *et al*., 2012) and adjusted to the average weight of mice. We culminated the experiment at 13 weeks of age of mice, which represents an early stage of disease based on the previous survival analysis in *Apc^Min/+^*mice that showed a drop in survival after 18-20 weeks of age(Curio *et al*, 2022). Food consumption, water intake and body weight were monitored weekly throughout the study. Animals were blind scored for the development of macroscopic signs of disease appearance (Supplementary Table 1). Prior to the commencing of the study, and at 4 and 7 weeks into the study, mice were put in metabolic cages overnight with *ad-libitum* access to food and water for the collection of urine and feces. Animals were singly housed to avoid cage-effects on microbial composition and metabolism throughout the study.

Tissue dissociation and flow cytometry, histology and immunohistochemistry and RNA extraction, proton nuclear magnetic resonance (^1^H NMR) spectroscopic analysis are described in Supplementary Information.

### RNA sequencing

RNA sequencing analysis was performed by Imperial BRC Genomics Facility (Imperial College London). Library was prepared using total RNA for cells and polyA mRNA for tissue samples. High-throughput sequencing was run in a HiSeq 4000 (Illumina) to obtain 75 bp sequencing paired-end reads. Sequencing data were mapped against a reference transcriptome with STAR alignment, followed by quantification using RSEM as detailed in alternate protocol 7 of the STAR (RRID:SCR_004463) software package(Dobin & Gingeras, 2015). Gene counts were generated using featureCounts(Liao *et al*, 2014) and subsequent analysis was performed using DESeq2 (RRID:SCR_000154) pipeline(Love *et al*, 2014) following the RNA-Seq workflow(Love *et al*, 2015). The DESeqDataSet object in cell RNA-seq data was built using the design formula ∼*Tyramine_treatment* (i.e. untreated, 1.2 mM tyramine or 1.6 mM tyramine) to determine the effect of tyramine treatment. For the mouse gut tissue RNA-seq data, genotype (*Apc^Min/+^* or WT) and treatment (tyramine or water) categories were combined in a new group, as we focused on the effect of tyramine treatment in *Apc^Min/+^* and WT groups, separately. The new group contained the following levels tyramine-treated *Apc^Min/+^*, water-treated *Apc^Min/+^*, tyramine-treated WT, and water-treated WT. The design formula used was as follows *∼ Gender +Parental + group.* Pre-filtering encompassed removing rows with less than 10 counts across all samples. We considered a gene to be differentially expressed if the false discovery rate (FDR) <0.05, and the absolute log2 fold change (LFC) >1 Gene ontology (GO) analysis was performed using goseq package as it takes into account gene length bias(Young *et al*, 2010). Gene Set Enrichment Analysis (GSEA) was applied to identify gene sets correlated with the phenotypic class of interest based on a ranked gene list(Subramanian *et al*, 2005). Gage package for gene set or pathway analysis was also used(Luo *et al*, 2009). Ingenuity Pathway analysis (IPA, RRID:SCR_008653) (Qiagen) was applied for pathway analysis. Graphs were plotted using ggplot (RRID:SCR_014601) package(Ginestet, 2011) in R.

### Statistical analysis

Statistical analysis was performed using Prism software (GraphPad, RRID:SCR_002798) and R (version 4.2.1). Data were assessed for normal distribution using the Kolmogorov-Smirnov normality test and the relevant statistic test was applied. No samples or animals were excluded from the analyses. Normally distributed data were analyzed by One-Way ANOVA with Dunnett’s or Tukey’s multiple comparisons test or unpaired t-test test as appropriate to the number of comparisons being made. Data that did not exhibit a normal distribution were analyzed using the nonparametric Kruskal–Wallis test with Dunn’s multiple comparisons test or Mann-Whitney test as appropriate to the number of comparisons being made. For grouped data, two-way ANOVA or mixed effects model followed by Tukey’s multiple comparisons test was used. Correlation analysis was performed using the Spearman correlation coefficient. Contingency table analysis using Fisher’s exact was applied. P < 0.05 was considered as statistically significant (*, p < 0.05, **, p < 0.01, ***, p < 0.001, ****, p < 0.0001).

## Acknowledgements

We thank L. Zarate-Garcia and J. Rowley from the Imperial College Flow Facility, L. Lawrence from the Imperial College Research Histology Facility, K. Petropoulou and R.D. Williams from Imperial Genomics Facility, and N. Rahman and A. Clear from BCI Pathology core services at Barts Cancer Institute for technical support. We thank Zhigang Liu, Yang Bi and Rachael Barry for help animal experiments. We thank Qianxin Wu, Toshiyasu Suzuki, and Evangelos Chandakas for critical discussions on the project. We want to thank CBS facility for their support.

## Author contributions

M.G. designed and performed experiments, analyzed data and wrote the manuscript. S.C designed and performed part of flow cytometry experiments, and analyzed data. S.S. analyzed RNA-seq data using GSEA pathway software analysis. M.E.B performed histological scoring for dysplasia. Y.W. supervised the RNA-seq study. N.G. supervised the flow cytometry study. N.J.G supervised the cell culture work. J.V.L obtained the funding, designed and supervised all the studies, analyzed data and wrote the manuscript. All authors edited and reviewed the manuscript.

## Data availability

The RNA-seq and NMR spectral data will be made available on National Center for Biotechnology Information (NCBI) and MetaboLights, respectively upon the acceptance of the manuscript.

